# PubData: search engine for bioinformatics databases worldwide

**DOI:** 10.1101/069575

**Authors:** Bohdan B. Khomtchouk, Kasra A. Vand, Thor Wahlestedt, Kelly Khomtchouk, Mohammed K. Sayed, Claes Wahlestedt

## Abstract

We propose a search engine and file retrieval system for all bioinformatics databases worldwide. PubData searches biomedical data in a user-friendly fashion similar to how PubMed searches biomedical literature. PubData is built on novel network programming, natural language processing, and artificial intelligence algorithms that can patch into the file transfer protocol servers of any user-specified bioinformatics database, query its contents, retrieve files for download, and adapt to the user’s search preferences.

PubData is hosted as a user-friendly, cross-platform graphical user interface program developed using PyQt: http://www.pubdata.bio. The methods are implemented in Python, and are available as part of the PubData project at: https://github.com/Bohdan-Khomtchouk/PubData.

## Introduction

If there was a data-oriented counterpart to PubMed Central (Roberts, 2001) in today’s ever-expanding data world, it would undoubtedly be PubData, a centralized repository dedicated to data in the life sciences. While biomedical literature is typically accessed via search engines such as PubMed or other software tools (Hokamp & Wolfe 2004, Fontaine et al. 2009, States et al. 2009, Lu 2011, Wang et al. 2014, Gou et al. 2015, Squizzato et al. 2015, bioCADDIE 2016, FORCE11 2016), the process of searching biomedical data often entails circuitously accessing various bioinformatics databases on a case-by-case basis, depending on the nature of the data sought by the user. Likewise, the search process often begins at the literature level, whereupon the user retrieves the respective open-access data only once the appropriate literary sources are located. This, in turn, impedes the efficiency of the data search process and prohibits the user from searching the other way around (e.g finding a list of papers that match the specifics of the data requested by the user).

Although biomedical data repositories such as the Gene Expression Omnibus (Edgar et al. 2002, Barrett et al. 2013) and the Sequence Read Archive (Kodama et al. 2012) offer a fairly comprehensive data search experience, they do not support the search of niche databases, which coincidentally comprise a significant majority of the ever-growing bioinformatics database ecosystem. Likewise, existing resources do not allow the user to search multiple databases simultaneously in a case-specific manner (e.g search concurrently in Uniprot, Ensembl, and the UCSC Genome Browser, but exclude other databases). In general, as the amount of data relevant to the biological sciences steadily grows and the number of bioinformatics databases progressively expands, a new initiative is imperative to establish a truly comprehensive electronic archive of data from peer-reviewed literature sources. This kind of resource must be made flexible in searching and intuitively navigating existing data repositories.

To this end, we propose PubData, currently offered as a user-friendly, cross-platform (Mac OS X, Windows, Linux) graphical user interface (GUI) search engine capable of accessing the file transfer protocol (FTP) servers of any bioinformatics database in the world and searching/retrieving their contents. Although the idea of the application of search engine technology to bioinformatics has a rich history (Liebel et al. 2004, Liebel et al. 2005, Marinescu et al. 2005, Morrison et al. 2005, Page 2005, Hearst et al. 2007, Lewis et al. 2012, Mandloi & Chakrabarti 2015, DeFreitas et al. 2016, Zhang et al. 2016), it has never been attempted at such broad scale. As such, we propose the first search engine designed to search data files in bioinformatics databases worldwide. We aim to provide the scientific community with the ability to conduct a “Google-style” search for biomedical data, thereby taking advantage of a data-oriented resource akin to the literature-oriented resource of PubMed.

PubData can remotely access, search, and retrieve files from the deeply nested directory trees of any major bioinformatics database via a local computer network. By assembling all major bioinformatics databases under the roof of one software program, PubData allows the user to avoid the unnecessary hassle and non-standardized complexities inherent to accessing databases one-by-one using an Internet browser. PubData allows a user to query multiple databases simultaneously for user-specified keywords (e.g human, cancer, transcriptome), and to manually add support for any additional bioinformatics databases as they come into existence. As such, PubData allows researchers to access, search, view, and download files from the FTP servers of any major bioinformatics database directly from one centralized location. By using only a GUI, PubData allows the user to simultaneously surf multiple bioinformatics FTP servers directly from the comfort of their local computer.

## Results & Discussion

PubData (Figure 1) has been written in the Python programming language (Python Software Foundation, 2016) and the user-interface has been designed using PyQt, which is the Python binding of the cross-platform GUI toolkit Qt (PyQt, 2016). In order to download or access biological files of interest, users typically connect to FTP servers. This could be very time-consuming and the method has inherent flaws. For instance, it is impossible to search files based on a property, or search concurrently in multiple databases.

**Figure 1.**
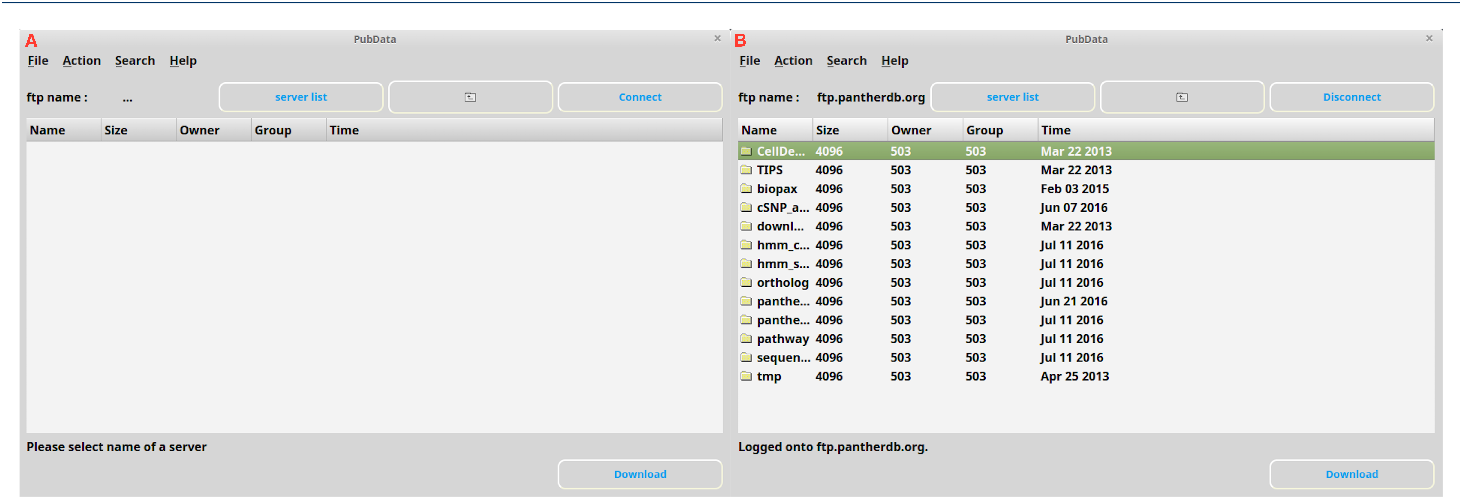
PubData user interface. (A) The PubData UI, from which the user can connect to, search, and download files from any number of bioinformatics databases provided by the server list widget. (B) Logged into PANTHER (Protein ANalysis THrough Evolutionary Relationships) Classification System database.

In PubData, we have resolved these issues. Now a user can search a specific keyword in several databases and see relevant results within seconds. PubData utilizes techniques in natural language processing (NLP) in order to retrieve information in an intuitive way. For instance, querying words that are semantically related (e.g. homo and human and mankind) leads to similar search results that are returned to a user within a short period of time. For implementation of the NLP module in PubData, we used the Natural Language Toolkit (NLTK, 2016).

The FTP (File Transfer Protocol) is a standard network protocol used to transfer computer files between a client and server on a computer network. Hence, in using FTP, there is no access to the entire directory tree. While traversing the directory tree on a server using FTP, users need to send a series of requests to a server (e.g. changing a path, downloading a file, retrieving the list of files in a directory, etc) and, as the size of the database gets larger, the transaction will inevitably take a lot of time (sometimes several hours). But since crawling/searching through a directory tree is the most time-consuming (and computationally expensive) procedure, we have designed PubData to perform this process locally as a background process rather than through FTP (Figure 2).

**Figure 2.**
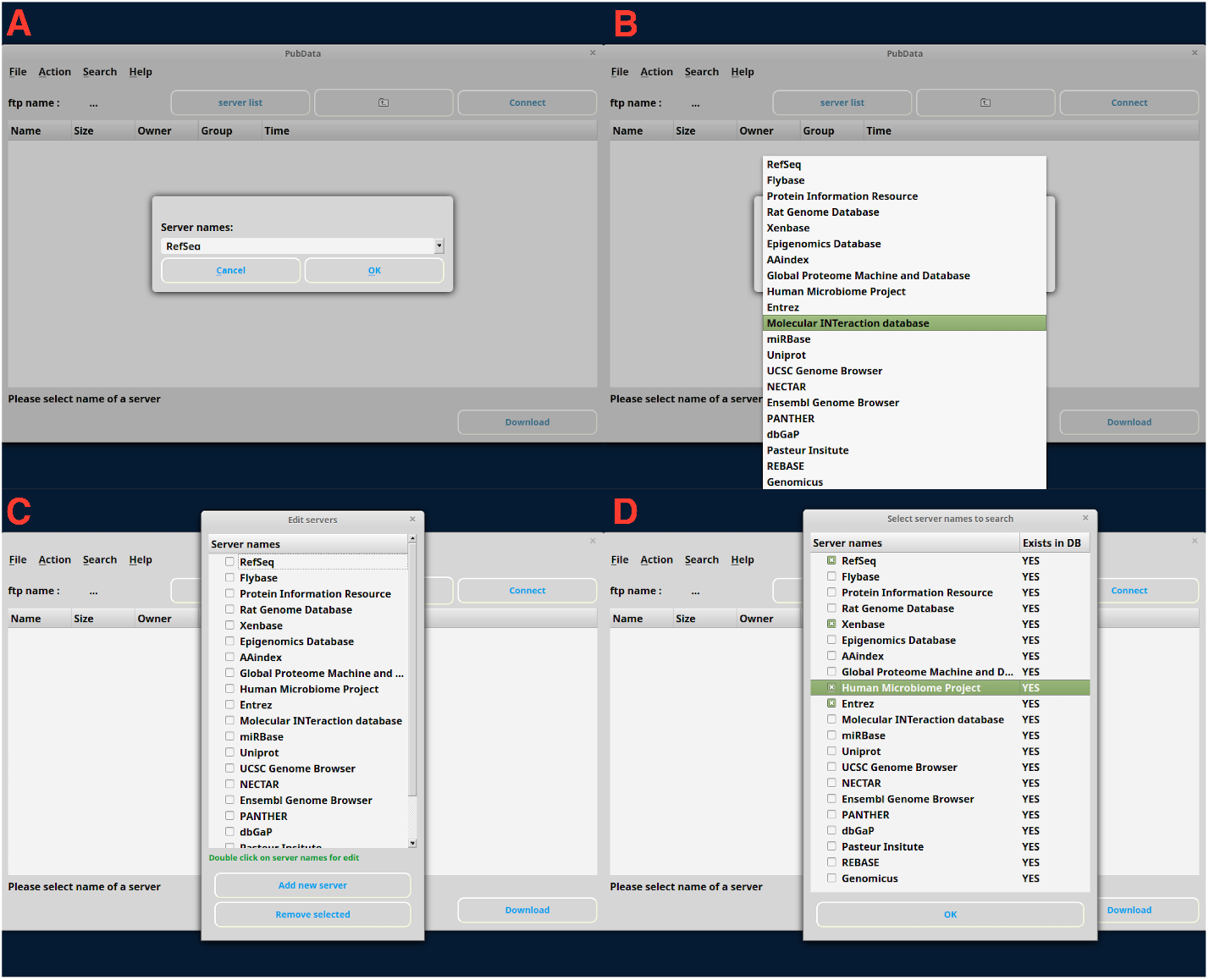
PubData search technology. (A) The user can choose a database to login to (e.g RefSeq) from a menu of bioinformatics databases (B). (C) If a user doesn’t see a specific database in the list, it is easy to manually insert a new database (convenient for recently published databases). (D) Searching multiple databases simultaneously via user-friendly checkboxes is useful for investigating search criteria concurrently across an array of databases.

To achieve this, we implemented a search module using the Breadth First Search (BFS) algorithm and a homebrewed FTP module, FTPWalker, and threaded this function through a parallelized and concurrent algorithm that divides the subdirectories within the root amongst multiple processes (based on available processors). Then, each process automatically divides the subdirectories between multiple threads that use FTPWalker in order to crawl through an FTP server’s entire directory tree. Since all we needed to deal with were filenames and their respective paths, we traversed all the FTP databases using the aforementioned method only once, and subsequently preserved the paths and filenames in a local SQLite database. The advantage of this approach is that at search time we can just search in this local database, and after extracting the relative paths we can simply access file directories by one single request, which makes all the computational processes extremely fast. The only drawback to this approach was that this method overlooked new files when FTP servers were updated. We circumvented this issue by providing the users with an update mechanism, so that users can manually update the databases whenever they like via a set of update buttons.

To efficiently crawl through an entire FTP server directory tree, a series of computational innovations in the form of parallelized algorithms were developed in PubData (e.g the FTPTraverse class and launcher.py). Although multithreading improves the performance of the application, there is no precise formula for calculating the algorithmic time complexity, since it depends on multiple factors. For instance, two threads can gain a speedup of up to 2X on-average, and four threads up to 3X. However, the potential speedup is bound to the available physical threads of the CPU and is dependent on balancing the computational workload. Specifically, we built a function called “find leading” that can find all directories and their respective subdirectories within the root directory. This function returns the leading directories when it encounters more than one directory with the corresponding path. Then the “main-walker” module divides the directories within root amongst the available processors by calling the “main run” function from the “traverse” module. Finally, this function borrows methods from the “find leading” function in order to find the subdirectories and partition them between threads (based on available ones) by calling the “traverse branch” function, which itself uses the “ftp-walker” module for traversing the FTP server. This module uses the BFS algorithm and Python’s “ftplib” module in order to be adapted for traversing an FTP directory tree. As such, performance gains depend mostly on the number of directories within the root and the number of subdirectories within them (i.e maximum depth). In general,executing two threads gives an increase in performance of up to 2X the original execution time. Using four threads, provides an increase of 2.5X to 3X the original execution time. Hence, if the number of CPUs is *C* and the regular execution time is *S*, the new execution time would approximately be *S/C* ×3 to *S/C* ×2.5

PubData provides users with efficient implementations of the following features:

- Selecting a server name and connecting to a corresponding FTP database in order to manually explore the database from root.
- Modification of the server list, include the ability to add new servers and deleting/editing the existing ones.
- Selecting an arbitrary number of local servers and searching among them.
- Automatically searching through all the servers.
- Suggesting the most frequently used words at search time, via a built-in recommender system (Figure 3).
- Match semantically related words via an NLTK general WordNet and a manual WordNet.
- Downloading any accessed files.
- Providing the metafile for each server, so that users can see the corresponding metafiles for each server after opening a specific path.

**Figure 3.**
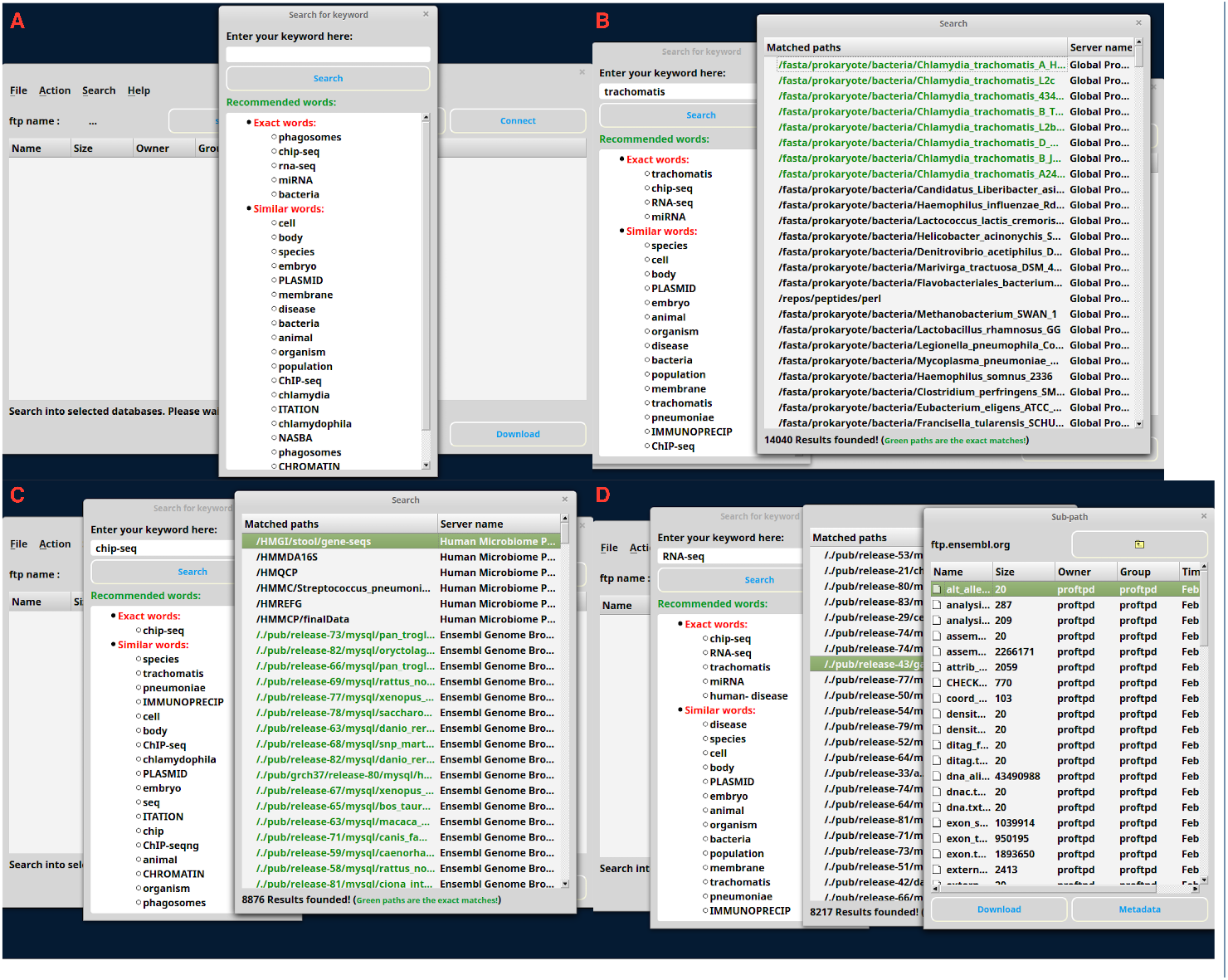
PubData recommender system. (A) Collaborative filtering recommender system based on previous search history. (B) Keyword search returns color-coded exact matches of the specific keyword to the specific file path. (C and D) Both exact matches and similar matches are returned by the search engine, allowing the user to pinpoint the full path destination to a desired group of files that meet the user’s search criteria.

Users will end up with a list of paths that contain files matching their queries, either semantically or literally (Figure 1). Then users can open a direct connection to a selected path and get access to a file directory on the server and download the files.

In general, to search a database, users enter a keyword and see two types of results:

- The keyword appears exactly in the file or directory names.
- The result is semantically and scientifically related to the keyword.

For semantic analysis, we used two kinds of WordNets. The first one is a general WordNet provided by the NLTK library, which only gives us some common relative results for a word. Here are some examples of synonyms given from the NLTK WordNet:

~~~
Sample input: “human”
Output: set([‘human being’, ‘homo’, ‘human’, ‘man’])

Sample input: “RNA”
Output: set([‘RNA’, ‘ribonucleic-acid’])
~~~

The NLTK corpora are general purpose and their coverage might not be adequate for an application in the realm of biology. Due this point and the fact that still there is no complete and free WordNet for biology, we created a new one. Here is the way that we created this biological WordNet:

We started by parsing two biological encyclopedias/dictionaries (Rittner & McCabe 2004, Singleton 2010). For all the words within these books, we extracted all the nouns from their corresponding description and then we refined those words by removing any general and irrelevant nouns. For example, here is a word alongside its relative nouns, as determined by PubData:

~~~
“adenylyl cyclase”: [“monophosphate”, “cytoplasm”, “signal”, “cAMP”, “molecule”, “adeno”, “enzyme”, “receptor”, “membrane”, “plasma”]
~~~

## Future Perspectives

We plan to continue improving the NLP aspects of PubData by adding more semantic analyses, as applied to query words, and improving the accuracy of the recommender system. We also plan to create a web-based version of PubData, allowing for platform-independent, web-browser accessibility for biologists.

## Conclusion

We provide access to a user-friendly graphical user interface program designed to access, search, retrieve, and download data files from any bioinformatics database in the world. We have gathered all the bioinformatics databases worldwide in PubData and let scientists search them quickly and efficiently. In addition, the use of semantic analysis has made the search engine retrieval system more intelligent and useful via a built-in recommender system. Since we realize that PubData is a community-driven project, we have made available the source code on GitHub with the vision that this encourages scientists and developers to help us improve the application. To encourage open-source contributions to PubData, we plan to determine future co-authorship on subsequent PubData publications by the number and quality of Github commits from the developer community.

## Competing interests

The authors declare that they have no competing interests.

## Author’s contribution

BBK conceived the study. BBK and KAV wrote the code. BBK, KAV, TW, KK, and MKS tested the code. TW created the website. CW provided project resources and participated in the management of the source code and its coordination. BBK wrote the paper. All authors read and approved the final manuscript.

## Acknowledgements

BBK wishes to acknowledge the financial support of the United States Department of Defense (DoD) through the National Defense Science and Engineering Graduate Fellowship (NDSEG) Program: this research was conducted with Government support under and awarded by DoD, Army Research Office (ARO), National Defense Science and Engineering Graduate (NDSEG) Fellowship, 32 CFR 168a.

